# A designed high-affinity peptide that hijacks microtubule-based transport

**DOI:** 10.1101/2020.06.17.157255

**Authors:** Jessica A. Cross, Magda S. Chegkazi, Roberto A. Steiner, Derek N. Woolfson, Mark P. Dodding

**Affiliations:** School of Chemistry, University of Bristol, Cantock’s Close, Bristol BS8 1TS, UK; School of Biochemistry, Faculty of Life Sciences, University of Bristol, University Walk, BS8 1TD, UK; Randall Centre of Cell and Molecular Biophysics, Faculty of Life Sciences and Medicine, King’s College London, London, UK; Bristol BioDesign Institute, University of Bristol, Life Sciences Building, Tyndall Avenue, Bristol BS8 1TQ, UK

## Abstract

Technologies that manipulate and augment the transport of vesicles and organelles by motor proteins along microtubules offer new routes to understanding its mechanistic basis, and could lead to therapeutics. Many cargoes for the kinesin-1 family of microtubule motors utilize adaptor proteins that harbor linear peptide motifs that are recognized by the tetratricopeptide repeats of kinesin light chains (KLC^TPR^s). These motifs bind with micromolar affinities at independent but overlapping sites. Here, we employ a fragment-linking peptide design strategy to generate an extended synthetic ligand (KinTag) with low nanomolar affinity for KLC^TPR^s. The X-ray crystal structure of the KLC^TPR^:KinTag complex demonstrates interactions as designed. Moreover, KinTag functions in cells to promote the transport of lysosomes with a high efficiency that correlates with its enhanced affinity. Together, these data demonstrate a new strategy for peptide design and its application to reveal that the more tightly a motor holds its cargo, the greater is the extent of cargo transport.

## INTRODUCTION

Up to 40% of protein-protein interactions (PPIs) are thought to be directed by the recognition of short linear peptides by larger protein domains (1, 2). A subset of these PPIs occur through the recognition of linear motifs by α-solenoid architectures such as armadillo repeat proteins (ArmRPs), HEAT proteins, and tetratricopeptide repeat (TPR) domains (3, 4). In these proteins, tandem arrays of α-helices are stacked together to form extended superhelical structures. This arrangement results in high surface-area-to-volume ratios compared with typical globular proteins and, thus, larger binding surfaces (5). TPR domains are important for a diverse range of PPIs in the cell including cell-cycle regulation, chaperone activity, transcription, protein translocation as well as intracellular transport and membrane trafficking (6). Specifically, TPR domain–peptide interactions play a critical role in cargo recognition by the microtubule motor kinesin-1 where the kinesin light chains TPR domains (hereafter denoted KLC^TPR^) recongize short unstructured peptides within cargo adaptor proteins, providing a direct link between motor protein and vesicular or organelle cargo to be transported (Figure 1a, reviewed in (7)). It is becoming clear generally that α-solenoid domain-peptide interactions are amenable to design, engineering and chemical manipulation, making them potential therapeutic targets for a range of diseases and the development of research tools (8-10).

**Fig. 1.**
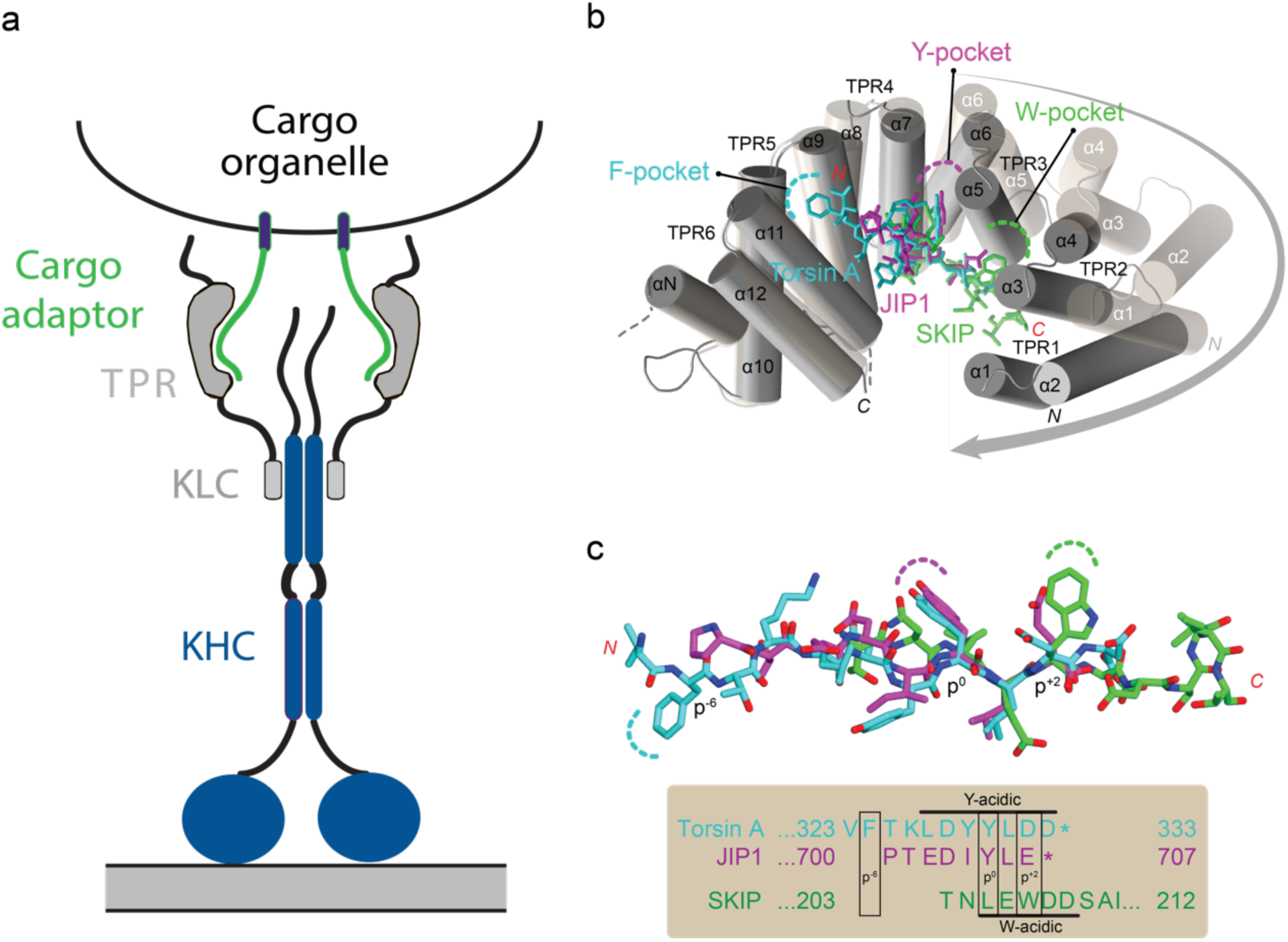
Kinesin-1 and recognition of cargo adaptor peptides. **(a)** Schematic of the kinesin-1 tetramer formed by two kinesin heavy chains (KHCs) and two kinesin light chains (KLCs). Cargo adaptor peptides are recognized by the tetratricopeptide repeat (TPR) domains of KLCs (KLC^TPR^s). **(b)** Cartoon representation of the cargo adaptor peptides of SKIP (green sticks, PDB code, 3ZFW), JIP1 (magenta, 6FUZ), and Torsin A (cyan, 6FV0) bound to the concave surface of KLC^TPR^ (solid gray helices). All peptides bind in the direction indicated by the red *N* and *C* letters. The KLC^TPR^ domain is formed of six helix-turn-helix TPR repeats (helices α1 – α12) and an additional non-TPR helix (αN) between TPR 5 and TPR 6. A flexible region between αN and α11 is indicated by the broken line. For simplicity, only one representative peptide-bound KLC^TPR^ is shown. The KLC^TPR^ domain in the peptide-free form (PDB code, 3NF1) is also shown as transparent cylinders with the arrow representing the domain closure elicited by peptide binding. Key binding pockets (F-, Y- and W-pockets) on KLC^TPR^ are highlighted. **(c)** Structure-based sequence alignment of the W-acidic motif of SKIP and the *C*-terminal Y-acidic motifs of JIP1 and Torsin A (asterisk indicates *C* terminus). Their color code is as in panel **b** with nitrogen and oxygen atoms in blue and red, respectively. The labels p^+2^, p^0^, and p^-6^ refer to the relative positions of peptide residues buried in the W-, Y-, and F-pockets of KLC^TPR^ with the Y residue of the Y-acidic motif as reference (p^0^).

Our own work and that of others has identified identified two distinct classes of peptide sequences that are recognised by KLC^TPR^: tryptophan-acidic (W-acidic) motifs found in the lysosomal cargo adaptor SKIP as well as many others feature a tryptophan residue flanked by aspartic or glutamic acids in the consensus L/M-D/E-W-D/E (7, 11-16); and Y-acidic motifs at the *C* termini of the axonal transport cargo adaptor JIP1 and Torsin A display a tyrosine residue flanked by acidic residues (17-20). In mammals there are four closely related KLC isoforms (KLC1 – 4). Whilst W-acidic motifs typically bind both KLC1^TPR^ and KLC2^TPR^, *C*-terminal Y-acidic motifs bind KLC1^TPR^ with a much higher affinity compared to KLC2^TPR^ (17, 19, 20). These motifs bind at distinct yet overlapping locations on the concave surface of the KLC^TPR^. Peptide binding increases the curvature of KLC^TPR^ through an induced-fit mechanism that highlights the adaptive plasticity of this domain as a platform for protein interactions (Figure 1b) (14, 17).

These natural motor-cargo adaptor interactions have µM affinities *in vitro* (14, 17, 19, 20). Previously, we have proposed that tighter interactions are achieved in the cell through avidity, as multiple copies of the adaptor peptides are expressed on the surface of organelles, they may be presented as a pair of motifs in adaptor proteins, and the kinesin-1 heterotetramer presents a pair of KLCs (Figure 1a) (7, 11, 21). Consistent with this proposition, fusion of multiple copies of W-acidic motifs in series to organelle associated proteins enhances transport compared with single motifs and with three motifs performing better than two motifs (22, 23). However, the hypothesis that adaptor-motor binding affinity *per se* is a limiting factor in transport is yet to be formally tested. We considered that one way to do this would be to design a ligand with higher affinity and ask how that affects transport.

More broadly, adaptable and synthetically accessible peptides are attractive candidates for disrupting PPIs as they mimic the natural interactions to compete for binding (24). However, protein-peptide interactions are often dynamic and weak due to the conformational lability of linear peptides. Thus, a challenge is to design peptide ligands that overcome this limitation by making additional or improved interactions with the target (25, 26).

We identified an opportunity to explore a new peptide design approach in the context of a structurally-defined transport system with a clear cellular functional readout, and at the same time, to ask a long standing question in kinesin biology. Here we describe a structure-guided, fragment-linking approach, which we call ‘*mash-up’* design, to deliver a high-affinity peptide ligand that targets the kinesin-1:cargo interface. This combines functional elements of different natural cargo-adaptor sequences into one peptide. Notably, it integrates the transport-activating capacity of W-acidic sequences with the extended binding interfaces from Y-acidic sequences. Our approach borrows from consensus design, which identifies key amino acids for a protein structure from multiple-sequence alignments (27, 28). However, it builds on this by incorporating offset fragments, analogous to fragment-based small-molecule drug design (29, 30). The *de novo* peptide, which we call KinTag, is effective in hijacking the transport function of endogenous kinesin motors in cells, and can deliver cargo into axons of primary neurons with a high efficiency that correlates with its enhanced binding affinity.

## RESULTS

### *Mash-up* design of peptide ligands for KLC TPR domains

The tryptophan and tyrosine residues in W-acidic and Y-acidic motifs, respectively, are essential for binding (14, 17-19). The X-ray crystal structures of cargo-adaptor peptides from SKIP, JIP1 and Torsin A complexed with KLC^TPR^ domains highlight how these residues interact with their receptor. In the KLC2^TPR^:SKIP complex, the central tryptophan (W, p^+2^ in Fig. 1c) residue occupies a hydrophobic pocket formed between TPRs 2 and 3 (the W-pocket) whilst the Y-acidic tyrosine (Y, p^0^) of both JIP1 and Torsin A bound to KLC1^TPR^ occupies a second pocket formed by repeats 3 and 4 (Y-pocket). The latter can also accommodate leucine or methionine residues often found at this position in W-acidic motifs. For Torsin A, an additional phenylalanine (F, p^-6^) occupies a third pocket (F-pocket) formed by TPRs 5 and 6 (Fig. 1b). Key to peptide stabilization within the concave surface of KLCs^TPR^ is an induced-fit closure of the receptors’ solenoid-shaped architecture that enables further hydrogen-bond formation.

A structure-guided overlay of SKIP, JIP1, and Torsin A KLC^TPR^-bound cargo adaptor peptides (Fig. 1c) suggested the possibility of simultaneously engaging the F-, Y- and W-pockets with a single peptide. Therefore, we sought to combine these different features of the natural sequences into a single synthetic-peptide ligand for KLC^TPR^ domains. To do this, we used a ‘*mash-up’* approach to design peptide sequences that extend the *N* terminus of the SKIP W-acidic motif with residues from the JIP1 (J-S peptides), or with residues from both JIP1 and Torsin A (T-J-S peptides). Since the p^-2^ leucine of SKIP and the p^0^ tyrosine from JIP1 and Torsin A bind to the same Y-pocket, two peptides each were designed for each of the J-S and T-J-S constructs (Table 1) (14, 17). The designs of these four peptides were made by solid-phase peptide synthesis with the 5-carboxytetramethylrhodamine (TAMRA) fluorophore appended *N* terminally. These were purified by high performance liquid chromatography and confirmed by MALDI-TOF mass spectrometry (Supplementary Fig.1, Supplementary Table 1).

**Table 1.**
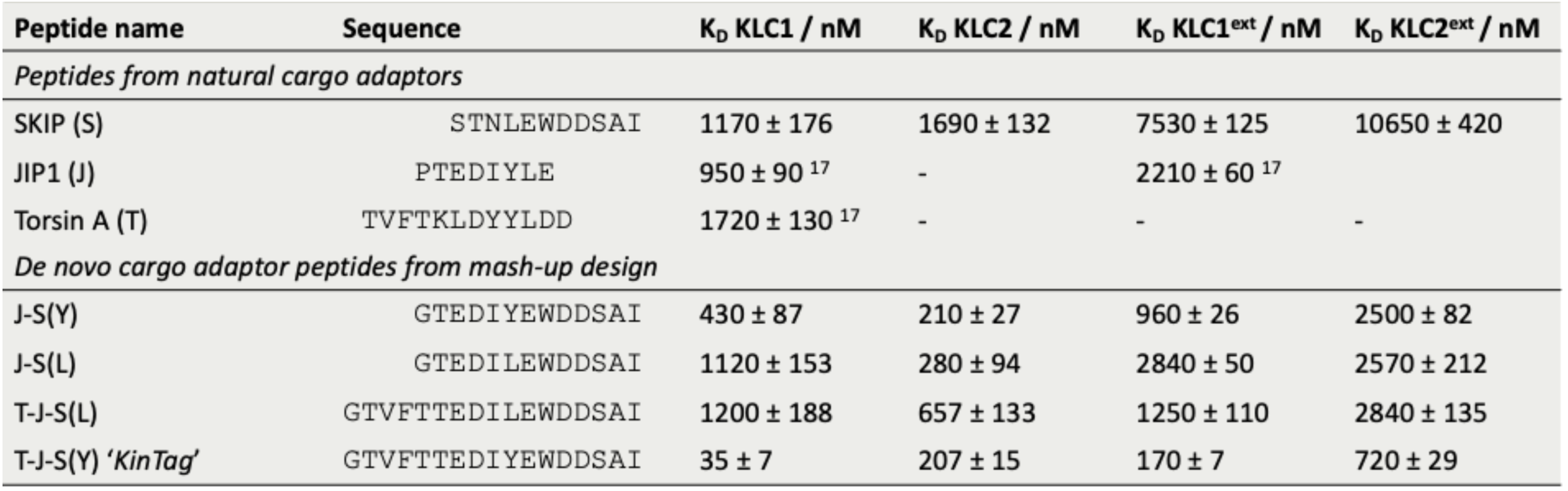
Sequences and *in vitro* binding affinities of natural and designed cargo adaptor peptides

### Synthetic peptides bind KLC^TPR^s *in vitro* with high affinities

The binding affinities of the designed peptides to KLC^TPR^ proteins comprising TPRs 1 – 6 were determined *in vitro* using fluorescence polarization (FP) assays. TAMRA-labelled peptides were incubated with increasing concentrations of KLC^TPR^ proteins in saturation-binding experiments(17) to obtain dissociation constants, *K*_D_, for both KLC1^TPR^ and KLC2^TPR^ (Fig. 2, Table 1). The designed peptides had increased affinities over the natural sequences for both isoforms. Peptides J-S(L) and J-S(Y) bound with moderately increased affinities, with the tyrosine variant as the tighter binder. The lower affinity of the peptides with p^0^ occupied by L, compared to Y, could be due to the loss of hydrogen bonds in the Y-pocket (Supplementary Fig. 3). Increasing the *N-* terminal extension in the T-J-S(Y) peptide led to a further increase in affinity, but this was not observed for the leucine variant. Thus, peptide T-J-S(Y) had the highest affinities for KLC^TPR^ with *K*_D_s of tens of nM to KLC1^TPR^ and hundreds of nM to KLC2^TPR^, Table 1. We dubbed this ‘KinTag’.

**Fig. 2.**
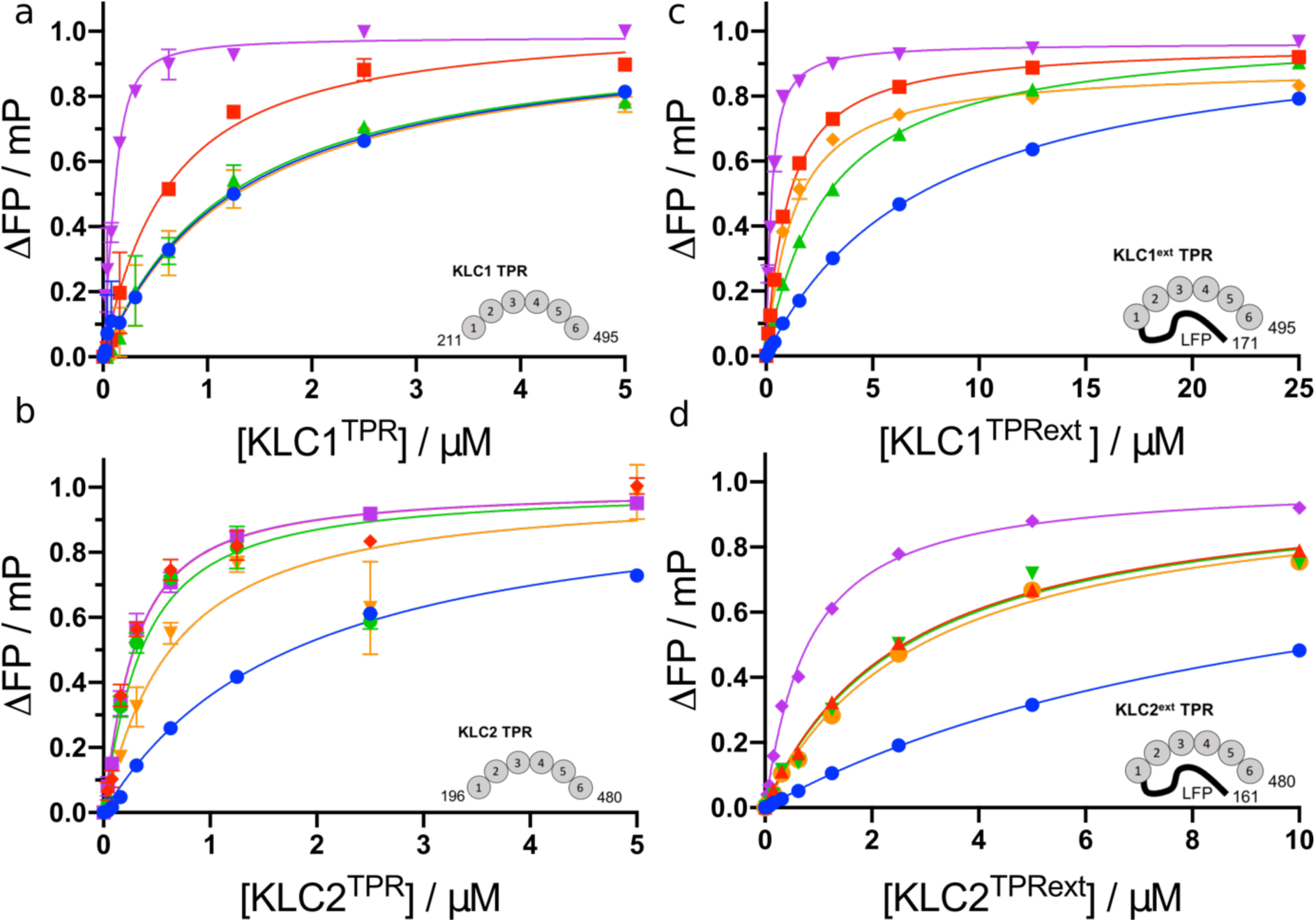
*In vitro* binding affinities of peptides for TPR domains. Fluorescence polarization assays for TAMRA-labelled peptides (150 nM) incubated with increasing concentrations of protein: **(a)** KLC1^TPR^, **(b)** KLC2^TPR^, **(c)** KLC1^TPRext^, and **(d)** KLC2^TPRext^. Key: natural SKIP, blue; J-S(Y), red; J-S(L), green; T-J-S(L), orange; and T-J-S(Y), KinTag, purple. Data were fitted to single-site binding models with polarization values of the peptide and buffer alone subtracted and values normalised to the calculated B_max_. Error bars indicate 1 SD from a minimum of 3 replicates.

**Fig. 3.**
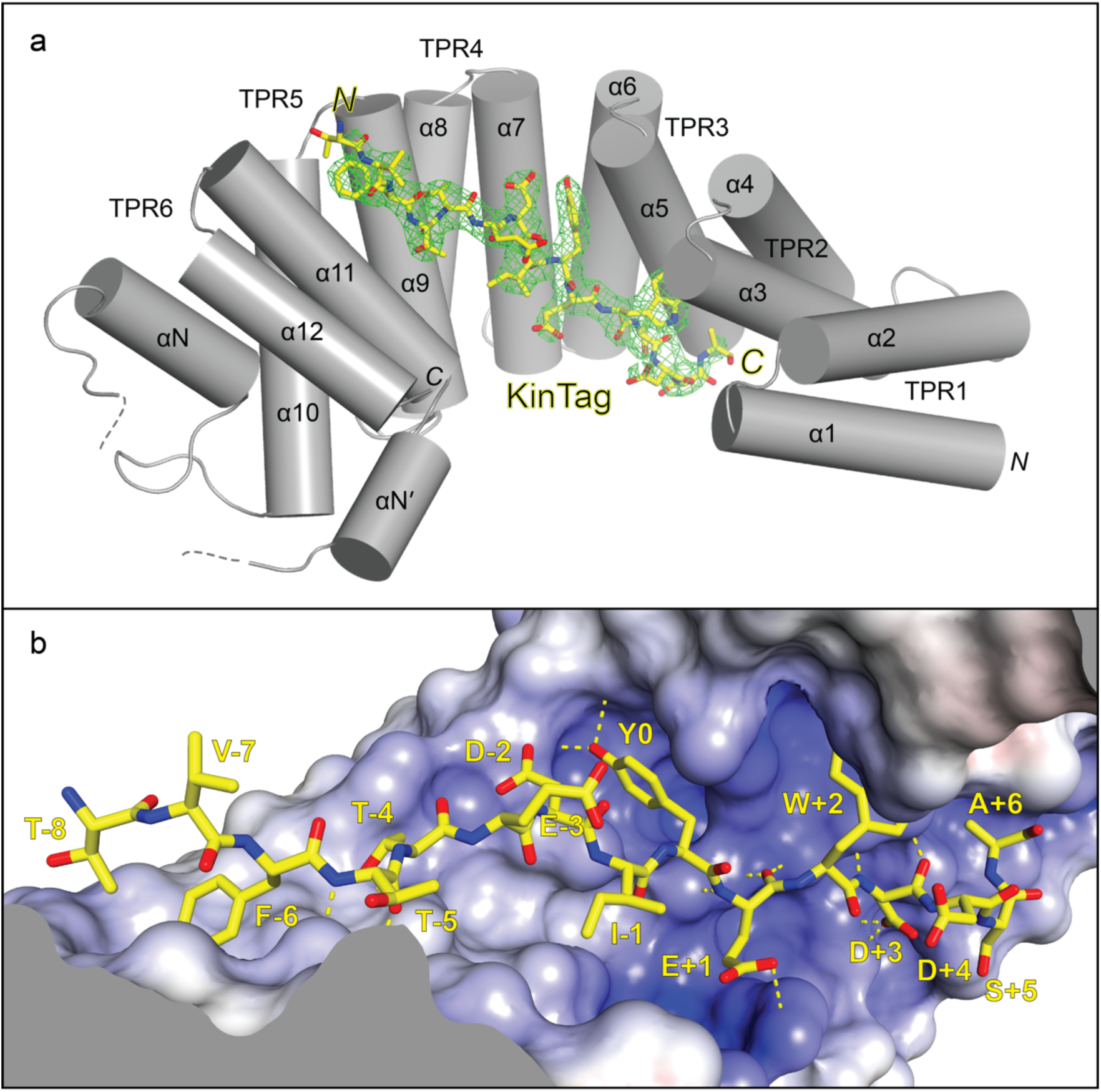
X-ray crystal structure of the KLC1^TPR^:KinTag complex. **(a)** Illustrated representation of the synthetic KinTag peptide (displayed as a yellow stick model accompanied by its *2mF*_*o*_*-DF*_*c*_ electron density in green contoured at the 1.0 σ level) bound to KLC1^TPR^. Helices α1 to α12 of the six TPR helix-turn-helix motifs are labelled. In addition to the non-TPR helix (αN) seen in previous structures another short helix (αN’) is revealed before α11. The flexible region between αN and αN’ is indicated with a broken line while the engineered disordered linker connecting α12 to the KinTag is not shown for clarity. Color-coded *N* and *C* labels indicate the *N* and *C* termini, respectively. Nitrogen and oxygen atoms are colored dark blue and red, respectively. **(b)** Sliced-surface view of KLC1^TPR^ with bound KinTag. The molecular surface is coloured according to its electrostatic potential with positive and negative potential in the +10*k*_B_T/e to -10*k*_B_T/e range shown in blue and red, respectively. Ordered KinTag residues are numbered with the central tyrosine residue as reference. Hydrogen bonds and salt bridges are shown as broken yellow lines.

### Designed peptides are sensitive to KLC^TPR^ autoinhibition

KLCs have a conserved autoinhibitory leucine-phenylalanine-proline-containing sequence (LFP-acidic) *N* terminal to their TPR domains (31). This regulatory element binds near TPR2 of KLC^TPR^, partly overlaps the W-acidic binding region, and forms part of a switch that triggers kinesin-1 conformational changes and activation (12, 31). In FP experiments, the presence of this region reduces the affinity of W-acidic peptides for the TPR domains, but not that of the Y-acidic peptide of JIP1, explaining why W-acidic peptides can initiate kinesin-1 activation but Y-acidic peptides do not (12, 31).

Consequently, this reduction in binding affinity can be interpreted as a proxy for the capacity to displace the inhibitory interaction (17, 31). Since a functional, transport-activating peptide would require this ability, we investigated the binding of the designed peptides to TPR constructs incorporating the conserved LFP-acidic region, KLC1^extTPR^ and KLC2^extTPR^ (Figs. 2c,d).

First, the designed peptides all had improved affinities over the natural sequences. KinTag bound to KLC1^extTPR^ with *K*_D_ = 170 nM, which is a 13-to 44-fold increase over the parent peptides (Table 1). Moreover, and importantly, the affinities of the designed peptides for KLC^extTPR^ were all reduced compared to those for KLC^TPR^. This demonstrates the sensitivity of these synthetic adaptor peptides to the presence of the LFP autoinhibitory motif, which we attribute to the W-acidic component of KinTag. In turn, we anticipate that these *de novo* cargo-adaptor peptides should retain the capacity to engage the LFP-acidic activation switch.

### The X-ray crystal structure of the KLC1^TPR^:KinTag complex reveals interactions as designed

To validate our mash-up design approach, we solved the X-ray crystal structure of the KLC1^TPR^: KinTag complex at the 2.85 Å resolution (Fig. 3 and Supplementary Table 2). In the crystallographic asymmetric unit, we found six copies of the KLC1^TPR^: KinTag complex arranged in three independent head-to-tail dimers (Supplemtary Fig. 2). The KinTag core sequence, VFTTEDIYEWDDS, is well defined in all complexes while the *N*-terminal glycine-threonine and the *C*-terminal alanine-isoleucine dipeptides are more flexible and partly visible only in some of the molecules. As designed, the KinTag peptide binds in an extended conformation within the concave groove of the TPR domain (Fig. 3a) with an interface area of 950 Å^2^. This represents a modest increase of approximately 24% compared to SKIP W-acidic (769 Å^2^) and just 7% over the rather extensive Torsin A Y-acidic interface (885 Å^2^). A number of hydrogen bonds and salt bridges stabilisze the peptide within the concave KLC^TPR^ surface together with electrostatic complementarity between the negatively charged peptide and its receptor (Fig. 3b, Supplementary Fig. 3). While all amino acids of KinTag contribute to the interface with KLC1^TPR^, it is the aromatic (F, Y, and W) residues that appear to play the key roles with 70%, 88%, and 80% of their accessible surfaces buried, respectively.

### KinTag efficiently hijacks intracellular transport

To validate the KinTag design functionally, we tested its capacity to hijack endogenous kinesin-1 to effect organelle transport in cells. Fusion proteins comprising the lysosome-associated membrane protein (LAMP1) and the green fluorescent protein (mGFP) with and without an intervening KinTag were transiently expressed in HeLa cells. A construct with the natural SKIP W-acidic sequence in place of KinTag was used as a control (23). As expected (32), the control LAMP1-mGFP fusion incorporated into lysosomes, and mostly clustered in the juxtanuclear region with some dispersed through the cytoplasm (Figs. 4a-c). Consistent with some limited capacity to promote transport, the single SKIP W-acidic sequence promoted measurable dispersion of lysosomes towards the cell periphery as shown by cumulative LAMP intensity (described in detail in (21, 33)). By contrast, KinTag promoted a dramatic dispersion of lysosomes with prominent accumulation of the fusion proteins at cell vertices. Notably, the extent of peripheral transport correlated with the binding affinities measured *in vitro*, with the high-affinity binder effecting a greater degree of lysosome dispersion (Figs. 4a-c).

**Fig. 4.**
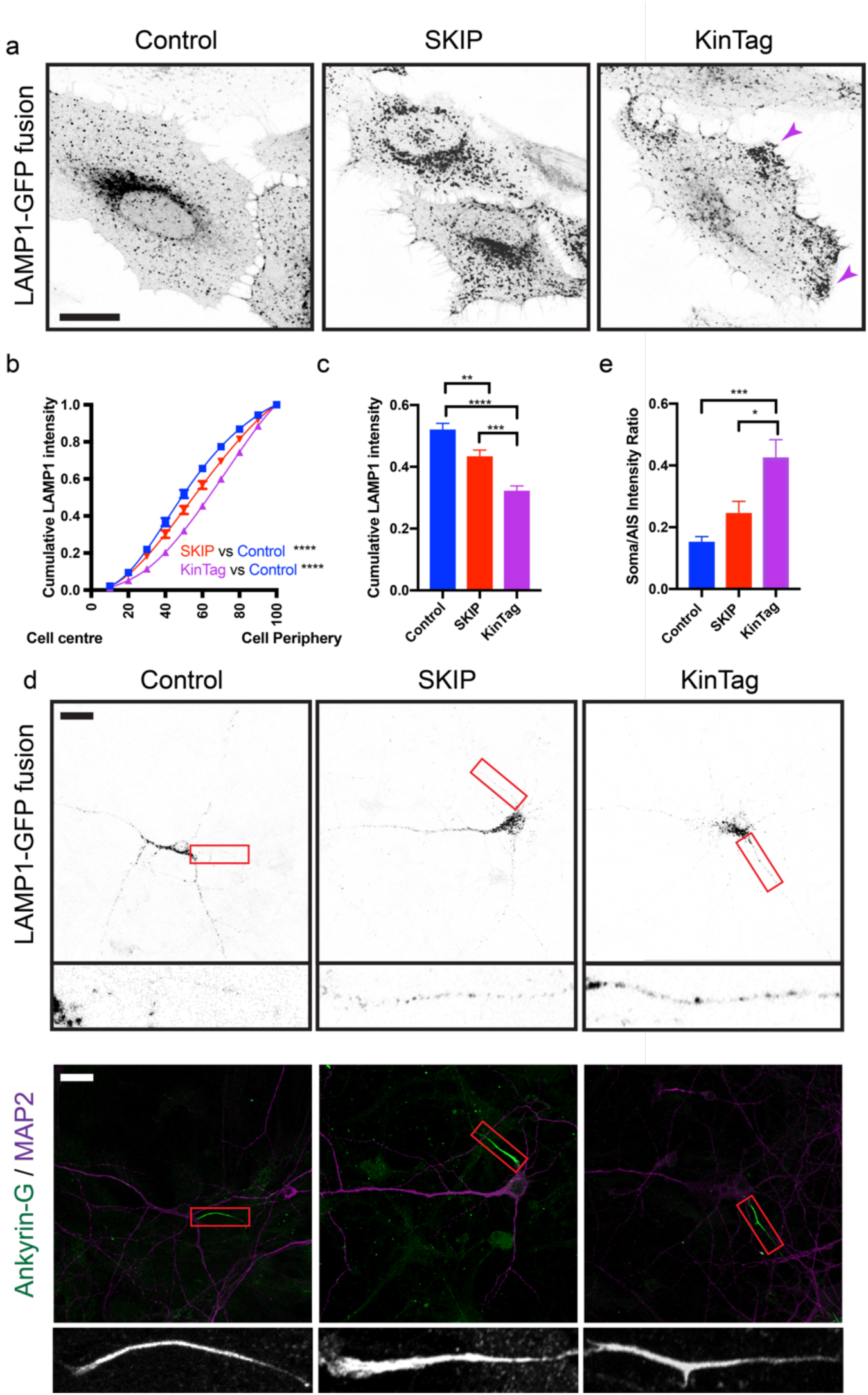
KinTag promotes more efficient lysosome transport than the natural W-acidic sequence. **(a)** Representative confocal fluorescence images of lysosome distribution in LAMP1-mGFP (control), LAMP1-SKIP-mGFP (SKIP) and LAMP1-KinTag-mGFP (KinTag) transfected HeLa cells. Scale bar is 20 µM. Purple arrows highlight fluorescence at cell vertices **(b)** Quantification of lysosome distribution by measuring cumulative fluorescence intensity from cell centre to cell periphery a minimum of 25 cells from 3 independent experiments. Fitted curves are compared using the extra sum of F-squares test (**c**) Comparision of cumulative LAMP1 intensity at the 50^th^ percentile. Error bars show S.E.M. and unpaired T-test is used for statistical analysis. **(d)** Representative confocal fluorescence images of lysosome distribution in LAMP1-mGFP (control), LAMP1-SKIP-mGFP (SKIP) and LAMP1-KinTag-mGFP (KinTag) in primary hippocampal neurons (inverted greyscale). MAP2 is shown in magenta **(e)** Quantification of ratio of fluorescence intensity in the soma and AIS in transfected neurons in a minimum of 12 cells from 3 independent experiments. Error bars show S.E.M. and unpaired T-test is used for statistical analysis. * P< 0.05, ** P< 0.01, *** P<0.001, **** P < 0.0001.

Finally, as kinesin-1 is known to be important for the transport of lysosomes into axons (22) we examined rat primary hippocampal neurons that were transfected with plasmids encoding LAMP1-mGFP, LAMP1-SKIP-mGFP and LAMP1-KinTag-mGFP fusion proteins. Entry of lysosomes into the axon was monitored by staining for ankyrin-G (an axon initial segment (AIS) marker) and quantified by measurement of the ratios of mean GFP fluorescence intensity in the AIS vs soma (Figs. 4d,e). Consistent with lysosome dispersion analysis in HeLa cells, KinTag promoted the robust transport of lysosomes into the axon (Figs. 4d,e). The SKIP W-acidic sequence was not as effective. For the first time, this demonstrates that the extent of kinesin-1-dependent axonal transport is positively correlated with the *in vitro* binding affinity of the interaction between a cargo adaptor peptide and the motor complex. This adds to examples of *in vitro* properties of *de novo* designed peptides translating to *in vivo* functions (34, 35). For the case described herein, it illustrates that *mash-up* design can deliver peptides that bind their targets with high affinity *in vitro*, and operate in cells to out-perform natural adaptor sequences and increase kinesin-1-dependent transport of a chosen cargo.

## Discussion

The recognition of client-peptide motifs by protein machineries is a ubiquitous feature of the dynamic cell. It enables different clients (*e*.*g*., integral membrane proteins) to be selected by conserved machineries (*e*.*g*., molecular motors or vesicle coats) to ensure the clients are directed to appropriate sites for their function, recycling, or degradation (7, 36). Here we have shown that this can be manipulated by combining structural insight from natural ligand-receptor interactions with synthetic peptide design.

We employ a novel *mash-up* design approach to combine binding features from natural sequences into a single, high-affinity peptide, KinTag, to hijack kinesin-1 dependent microtubule transport. At a molecular level, our design approach is validated by X-ray crystallography, and its functional utility is demonstrated in lysosome transport assays. For the first time, this reveals that protein-peptide binding affinities measured *in vitro* correlate with the extent of cargo transport in the cell. Our study opens the door to manipulating similar interactions throughout transport and trafficking networks, where similar unitarily low affinity natural interactions are typically supported through co-operativity, avidity and coincidence detection, and thus may be amenable to augmentation through the design of high-affinity peptide ligands.

More broadly, *mash-up* design provides a strategy for developing high-affinity peptide ligands for protein surfaces. The structure of the KLC1^TPR^:KinTag complex confirms that both general design features (*i*.*e*., shape and charge complementarity between the peptide and the protein) and sequence-specific design features (*i*.*e*., large side chains of the peptide bind into designated pockets of the protein) are realized. This design approach can yield an order of magnitude or more increase in binding affinity without dramtically increasing the interface area, which highlights the utility of fragment linking in peptide design, akin to that used with small-molecule pharmacophores (29, 30). Crucially, we have been able to incorporate a key function of a natural ligand—namely, operating the KLC autoinhibitory switch—(12, 31) demonstrating that complex functionality can be retained from peptide fragments.

Finally, we have shown that the KLC^TPR^s have a have a much broader capacity for selective motif recognition than is currently appreciated and are, in principle, capable of interacting with their cargo adaptor ligands over a much wider range of binding affinities. It seems likely that some natural ligands may also incorporate elements of the Y-acidic and W-acidic consensus, or perhaps utilize other pockets within the groove. Nonetheless, our data confirm the key role and intrinsic sufficiency of TPR groove binding peptides for motor recruitment and activation and through comparison of *in vitro* measured binding affinities with the extent of cargo transport, we have demonstrated, for the first time, that these are positively correlated. Put another way, we have established a simple concept: the more tightly a motor holds onto its cargo adaptor, the more efficiently the cargo is transported. These findings should allow a toolkit of peptide ligands, across a range of affinities, to be developed as transport tags within cells. This will allow investigation of the relationship between *in vitro* binding affinities and binding kinetics, with the cellular mechanics of vesicle and organelle transport.

We envisage wider applications of KinTag and similarly derived peptides in the orchestration of transport of organelles, and other cargoes, to investigate the functional roles of positioning within the cell, building on the the strategy of fusing tandem and triplicate motifs to transmembrane proteins previously explored Farias *et al*. (22), but now using a single, structurally and biophysically defined, short sequence. Moreover, it should be possible to augment binding affinity in existing motifs through amino-acid substitutions to explore the role of manipulating relative motor activity in systems where opposite polarity motors bind the same cargo or cargo adaptor (37). It is conceivable that the design of relatively low molecular weight, high-affinity, peptide or peptidomimetic ligands for kinesin-1 could be useful in the spatial (*e*.*g*., axon specific) targeting of exogenous therapeutic cargoes for the treatment of neurological disease.

## ACKNOWLEDGEMENTS

J.A.C is supported by the EPSRC Bristol Centre for Doctoral Training in Chemical Synthesis. M.P.D. is a Lister Institute for Preventative Medicine Research Prize Fellow and work in his lab is supported by BBSRC grant BB/S000917/1. D.N.W. held a Royal Society Wolfson Research Merit Award (WM140008). M.S.C. and R.A.S. are supported by BBSRC grant BB/S000828/1. We also thank the University of Bristol School of Chemistry Mass Spectrometry Facility for use of the EPSRC-funded Bruker Ultraflex MALDI-TOF/TOF instrument (EP/K03927X/1), and the BBSRC/EPSRC-funded BrisSynBio (BB/L01386×1) for access to its peptide synthesizers and plate reader. The X-ray crystallographic work was conducted at beamline I03 of Diamond Light Source. The authors also thank Kirsty McMillan for donation of rat hippocampal neurons and antibodies and assistance with experimental protocols.

## AUTHOR CONTRIBUTION

M.P.D., D.N.W and J.A.C. conceived the project and designed the peptides. J.A.C. synthesised the peptides and completed biophysical and cellular experiments. R.A.S. and M.S.C. designed the construct used for crystallographic studies and solved the X-ray crystal structure of the complex. The manuscript was written by J.A.C., M.P.D., D.N.W. and R.A.S. and commented on by all authors.

