## Supplementary Material for "A designed high-affinity peptide that hijacks microtubule-based transport"

* To whom correspondance should be addressed:

;

#### Materials and Methods

Solid phase peptide synthesis

Fmoc protected amino acids, dimethylformamide (DMF) and 6-Chloro-1-hydroxybenzotriazole (Cl-HOBt) were purchased from Cambridge Reagents. Rink amide supporting resin (100-200 mesh) was purchased from Novabiochem®. N, N′-Diisopropylcarbodiimide (DIC) was purchased from Carbosynth, morpholine from Merck Millipore, formic acid, Triisopropyl silane (TIPS), and Trifluoroacetic acid (TFA) from Acros Organics, pyridine and dichloromethane (DCM) from Fisher, acetic anhydride from BDH Laboratories, diethyl ether from Honeywell, and HPLC-grade acetonitrile (MeCN) from VWR Chemicals. 5-Carboxytetramethylrhodamine (TAMRA) was purchased from Novabiochem®, (1-[Bis(dimethylamino)methylene]-1H-1,2,3-triazolo[4,5-b]pyridinium 3-oxide hexafluorophosphate (HATU) and N,N-Diisopropylethylamine (DIPEA) were purchased from Sigma Alrdrich. All chemicals were used as supplied. Synthesis-grade ultra-pure DMF (Cambridge Reagents) was used exclusively during peptide synthesis. All structures included in the manuscript were created using educational-use PyMOL software (https://www.pymol.org/). Exact peptide masses were calculated using Peptide Synthetic’s online Peptide Mass Calculator tool (http://www.peptidesynthetics.co.uk/tools/)

Standard Fmoc solid-phase peptide synthesis was performed on a 0.1 mM scale using CEM (Buckingham, UK) Liberty Blue automated peptide synthesis apparatus with inline UV monitoring. Activation was achieved with DIC/Cl-HOBt. Fmoc deprotection was performed with 20% v/v morpholine/DMF with addition of 5% formic acid to prevent aspartimide formation. Double couplings were used for β-branched residues and the subsequent amino acid. All peptides were synthesised from C to N terminus as the C-terminal amide on Rink amide resin. TAMRA (0.1 mM, 2 eq.), HATU (0.095 mM, 1.9 eq.) and DIPEA (0.225 mM, 4.5 eq) in DMF (3 mL) were added to DMF washed peptide resin (0.05 mM) with agitation for 3 hours. Resin was washed with 20% piperidine in DMF (5 mL) for 2 x 30 minutes to remove any excess dye. All manipulations were carried out under foil to exclude light. Peptides were cleaved from the solid support by addition of TFA (9.5 mL), TIPS (0.25 mL) and water (0.25 mL) for 3 hours with shaking at rt. The cleavage solution was reduced to approximately 1 mL under a flow of nitrogen. Crude peptide was precipitated upon addition of ice-cold diethyl ether (40 mL) and recovered via centrifugation. The resulting precipitant was dissolved in 1:1 acetonitrile and water (≈ 15 mL) and lyophilised to yield crude peptide as a solid. Peptides were purified by reverse phase HPLC on a Phenomenex (Macclesfield, UK) Luna C18 stationary phase column (150 x 10 mm, 5 μM particle size, 100 A pore size) using a preparative JASCO HPLC system. A linear gradient of 10-60 % ammonium bicarbonate (25 mM) and water was typically applied over 30 minutes with gradient adjusted for each peptide to optimise separation. Chromatograms were monitored at wavelengths of 220 and 280 nm. The identities of the peptides were confirmed using MALDI-TOF mass spectrometry using a Bruker ultrafleXtreme II instrument in reflector mode. Peptides were spotted on a groundsteel target plate using dihydroxybenzoic acid as the matrix. Masses quoted are for the monoisotopic mass as the singly protonated species. Masses were measured to 0.1% accuracy. Peptide purities were determined using a JASCO analytical HPLC system, fitted with a reverse-phase Kinetex® C18 analytical column (Phenomenex, 5 μm particle size, 100 Å pore size, 100 x 4.6 mm). Fractions containing pure peptide were pooled and lyophilised. Peptides were dissolved in buffer (25 mM pH 7.4 HEPES buffer with 150mM NaCl, 5mM BME) and their concentrations determined by UV-Vis on a ThermoScientific (Hemel Hemstead, UK) Nanodrop 2000 spectrophotometer using measurement of UV absorbance at 555 nm ($\varepsilon_{555}$(TAMRA) = 85 000 mol$^{-1}$ cm$^{-1}$).

Plasmids

The TPR domain of KLC1 alone (KLC1$^{TPR}$, amino acids A211–S495), a longer version containing the conserved LFP motif (KLC1extTPR, K173–S495)), and equivalent constructs for KLC2 (KLC2$^{TPR}$, A196–S480 and KLC2$^{extTPR}$, K161–S480) cloned in the bacterial expression vector pET28His-Thrombin have been described previously.(1, 2) A codon-optimized DNA sequence encoding the TPR domain of mouse KLC1 (residues 205 – 496, Uniprot Q5UE59, KLC1$^{TPR}$) fused N-terminally to the KinTag peptide via via a (Thr-Gly-Ser)$_{10}$-Gly flexible connector was purchased from Genscript and subcloned between the NdeI/XhoI sites of a pET28 vector (Novagen). This strategy allowed for the expression of a chimeric protein KLC1$^{TPR}$-(TGS)$_{10}$-G-KinTag (KLC1$^{TPR}$:KinTag) bearing a thrombin-cleavable N-terminal hexa-histidine tag. A plasmid for LAMP1-mGFP was obtained from Addgene, plasmid number 34831(3), and SKIP or Kin-Tag sequences followed by a (TGS)$_2$ linker cloned between LAMP1 and mGFP in Sal1-BamH1 restriction enzyme sites by annealing of the following oligonucleotides:

SKIPFor
TCGACGGCAGCACCAACCTGGAGTGGGATGATAGCGCGATTACCGGCAGCAC
TGGATCAG
SKIPRev
GCCGTCGTGGTTGGACCTCACCCTACTATCGCGCTAATGGCCGTCGTGACCTA
GTCCTAG
KinTagFor:
TCGACGGCACCGTGTTTACCACCGAAGATATCTATGAATGGGATGATAGCGCG
ATTACCGGCAGCACTGGATCAG
KinTagRev:
GCCGTGGCACAAATGGTGGCTTCTATAGATACTTACCCTACTATCGCGCTAATG
GCCGTCGTGACCTAGTCCTAG

Protein expression

Single bacterial colonies of E.coli BL21(DE3) cells were picked and grown overnight in 5 ml LB supplemented with kanamycin (50 µg/ml) at 37°C, 180 rpm. Small scale overnight bacterial cultures (5 ml) were used to inoculate larger cultures (1 L) which were incubated at 37°C until they reached an OD600 of 0.8. Protein expression was induced by the addition of 300 µM Isopropyl β-D-1-thiogalactopyranoside (IPTG) and the cultures incubated at 16°C, 180 rpm for 16 hours. Cells were harvested by centrifugation at 4000 x g for 30 minutes at 4°C and resuspended in 35 ml lysis buffer (25 mM HEPES pH 7.4, 500 mM NaCl, 20mM imidazole, 5mM β-mercaptoethanol supplemented with protease inhibitor cocktail (Roche)). Cell lysis was accomplished by sonication for 3 minutes (Pulses of 5 seconds on, 10 seconds off, 70% amplitude).

Insoluble material was sedimented by centrifugation at 13 000 rpm for 30 minutes at 4°C and the supernatant loaded onto a His-trap HP column (GE Healthcare) preequilibrated with lysis buffer. The protein was eluted with an imidazole gradient (20 mM to 500 mM Imidazole) over 100 ml and fractions containing the target protein identified by SDS polyacrylamide gel electrophoresis. Protein was applied to a HiLoad 16⁄600 Superdex 75 prep grade column (GE Healthcare) preequilibrated with sample buffer (25 mM HEPES pH 7.4, 500mM NaCl, 5mM β- mercaptoethanol) and 2 ml fractions collected using an AKTA prime plus. Fractions containing pure protein were identified by SDS polyacrylamide gel electrophoresis, pooled and concentrated using Vivaspin® sample concentrators (Sigma Aldrich). Protein concentration was determined using a UV/Vis spectrometer. For crystallization, the KLC1$^{TPR}$:KinTag sample eluted from the immobilized metal affinity chromatography step was buffer exchanged in lysis buffer and was further incubated overnight at room temperature with thrombin conjugated beads (Thrombin CleanCleave Kit, Sigma-Aldrich). The protein-bead mixture was filtered using a gravitational column and non-tagged protein was collected utilizing reverse IMAC on a HisTrap HP column, followed by size exclusion chromatography on a HiLoad 16/600 Superdex 75 column (GE Healthcare) pre-equilibrated with 25 mM HEPES pH 7.5, 150 mM NaCl and 5mM *β*-mercaptoethanol.

Fluorescence anisotropy

TAMRA conjugated peptides were diluted to 150 nM and incubated with increasing concentrations of the protein of choice (typically 0 – 10 µM) in assay buffer (25mM HEPES pH 7.4, 5 mM β-mercaptoethanol and 150mM NaCl) using an EpMotion liquid handling robot (Eppendorf). Measurements were performed on a CLARIOstar (BMG Labtech) microplate reader at room temperature. Data analysis was performed using the Prism (GraphPad Software Inc., San Diego CA, USA) package. Data was fitted to a quadratic one-site tight binding equation to account to calculate a value for $K_d$ accounting for ligand depletion (Equation 1).(4)

$$\frac{[RL]}{[L]_{Tot}} = \frac{([R]+[L]+K_d)-\sqrt{([R]+[L]+K_d)^2-4[R][L]}}{2[L]}$$ **Equation 1**

Crystallography

Crystallization of KLC1$^{TPR}$:KinTag was performed by the vapor diffusion method using the sitting drop setup. More specifically, purified untagged KLC1$^{TPR}$:KinTag was concentrated to ~8 mg/ml using Vivaspin centrifugal concentrators (Merck) and crystallization trays were set up with the aid of the Mosquito liquid handler (TTP LabTech) in 96-well Swissci MRC plates (Molecular Dimensions). Crystals developed after two days at 19 ˚C, in 0.1 M MIB buffer (sodium malonate dibasic monohydrate, imidazole and boric acid) pH 9.0 and 25% (w/v) PEG 1500 (Pact Premier screen, Molecular Dimensions) using a 2:1 protein:precipitant ratio in 400nl drops. For data collection these crystals were cryo-protected using the reservoir conditions spiked with 30% (v/v) ethylene glycol. A complete dataset at the 2.85 Å resolution was collected at the I03 beam line of Diamond Light Source (Didcot, United Kingdom). Data processing was performed using the *xia2* pipeline(5, 6) and the crystal structure was solved by molecular replacement using *Phaser(7)* using an ensemble composed of the KLC1$^{TPR}$ domains from 3NF1(8) and 6FV0(2) as starting models. Model refinement was carried out with the *Refmac5* package of the *CCP4* software suite.(9) Model building was performed with *COOT.(10)* A summary of data collection and refinement statistics are shown in Table S2. Structural images were prepared with PyMol (Schrödinger).

<u>Transfection of HeLa cells</u>

HeLa cells were maintained in high glucose Dulbecco's Modified Eagle's Medium (Gibco Invitrogen) with 10 % (v/v) foetal calf serum (Sigma-Aldrich) and 5 % penicillin/streptomycin (PAA) (herein referred to as DMEM) at 37 °C and 5 % $CO_2$. For transfection, cells were seeded in 6-well plates on fibronectin coated 13 mm coverslips at a density of 1 x $10^5$ cells per well and incubated at 37 °C and 5 % $CO_2$ for 16 h prior to transfection. Cells were transfected with 0.4 µg DNA using Effectene transfection reagent according to the manufacturer's instructions (Qiagen). After transfection cells were incubated at 37 °C, 5 % $CO_2$ for 16 hours. Cells were fixed by addition of 4% paraformaldehyde in PBS at room temperature for 10 minutes (2 ml per well for 6-well plate) and washed 3 x with PBS. Confocal images were collected using a Leica SP5II system with a 40× or 60× objective running Leica LAS X and are presented as maximum intensity projections. Figures were assembled using ImageJ in conjunction with Inkscape.

<u>Lysosome distribution analysis in HeLa cells</u>

To quantify lysosome distribution in an unbiased fashion, widefield images of cells were acquired at 40× magnification. As described previously, the cell perimeter was defined by thresholding equivalent saturated images and the cell area was scaled in 10% decrements using ImageJ. After background subtraction, cumulative integrated LAMP1 intensity (relative to the whole cell) was then plotted for increasing incremental deciles.**(11)** Data points are from a minimum of 25 cells and are representative of at least 3 independent experiments. For analysis of the resulting data, the nonlinear regression function in Graphpad Prism was used to fit a centred 6th order polynomial. To compare models so assess the statistical significance of differences in distribution profiles, the extra sum of F-squares test was applied. P-values for particular comparisons are indicated on the graphs. In addition, the cumulative integrated LAMP1 intensity in the central 50% of the cell plotted for the different constructs and compared using an unpaired T-test function in Graphpad Prism.

<u>Transfection of Neuronal cells</u>

Primary hippocampal neuron cultures were prepared from embryonic day E18 Wistar rats as previously described.(12) Dissociated neurons were grown on 22 mm glass coverslips coated in poly-L-lysine (1 mg/mL, Sigma). Cells were plated at 150,000 cells/coverslip in 2 ml plating medium (Neurobasal (Gibco) supplemented with 5 % horse serum (Sigma); 1 % GlutaMAX (Gibco); and 2 % B27 (Gibco)). Media was changed to feeding medium (Neurobasal (Gibco) supplemented with 0.4 % GlutaMAX (Gibco); and 2 % B27 (Gibco)) 2 h after plating. Cells were fed with an additional 1 ml of feeding medium 7 days after plating. Cells were cultured in a humidified incubator at 37 °C and 5 % $CO_2$. Animal care and procedures were carried out in accordance with the UK Animals Scientific Procedures Act (1986) and University of Bristol guidelines.

Transfection of neuronal cultures was carried out at DIV 12 using Lipofectamine LTX (Invitrogen) according to the manufacturer's instructions. Cells were left for 48 h before fixation with 4 % (v/v) paraformaldehyde in PBS at room temperature for 15 minutes (2 ml per well for 6-well plate) and washed 3 x with PBS. Paraformaldehyde was quenched with the addition of 100 mM glycine. For immunofluorescence mouse anti-Ankyrin-G was from NeuroMab (clone N106/3) and Goat anti-Mouse Secondary Antibody, Alexa Fluor 568 (Thermo Fisher). Confocal images were collected using a Leica SP5II system with a 40× or 60× objective running Leica LAS X and are presented as maximum intensity projections. Figures were assembled using ImageJ in conjunction with Adobe Photoshop and Illustrator.

Lysosome distribution analysis in neurons

To quantify lysosome distribution in an unbiased fashion, images of cells were acquired at 40× magnification. The soma perimeter was defined by thresholding equivalent saturated images. The axonal initial segment (AIS) was defined by thresholding equivalent images stained for ankyrin G. After background subtraction, mean LAMP1 intensity for the two areas was plotted. Data points are from a minimum of 12 cells and are representative of at least 3 independent experiments. To compare data sets we used an unpaired T-test function in Graphpad Prism.

#### Supplementary Figures and Tables

| **Supplementary Table 1 \| Masses of synthetic peptides** | | | |
|---|---|---|---|
| **Peptide name** | **Sequence** | **Expected mass** | **Observed mass** |
| *Peptides from natural cargo adaptors* | | | |
| SKIP | GSTNLEWDDSAI | 1718.77 | 1719.106 |
| JIP1 | GPTEDIYLE | 1447.54 | 1448.546 |
| *De novo cargo adaptor peptides from mash-up design* | | | |
| J-S(Y) | GTEDIYEWDDSAI | 1924.96 | 1926.784 |
| J-S(L) | GTEDILEWDDSAI | 1874.95 | 1876.147 |
| T-J-S(L) | GTVFTTEDILEWDDSAI | 2323.46 | 2325.057 |
| T-J-S(Y) ‘KinTag’ | GTVFTTEDIYEWDDSAI | 2373.48 | 2374.887 |

### SKIP

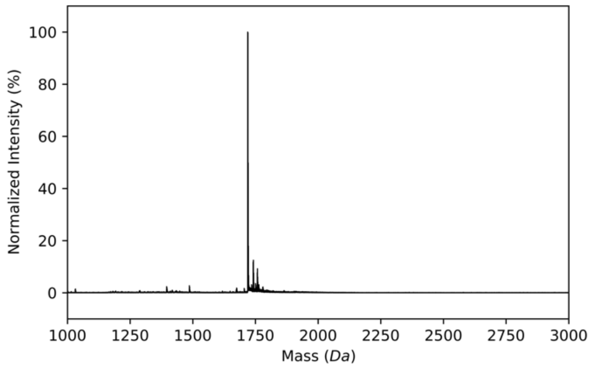


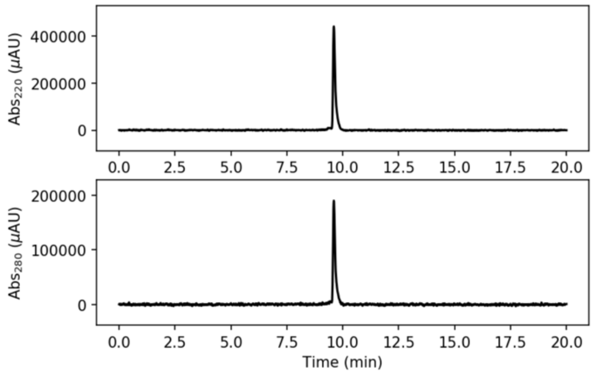


### JIP1

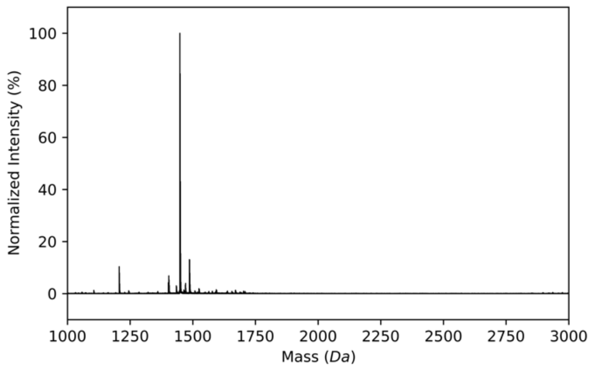


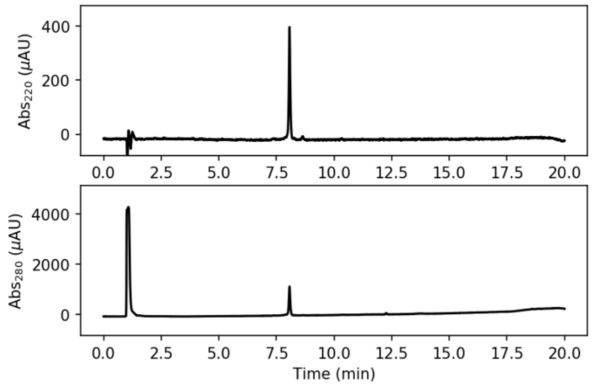


# J-S(Y)

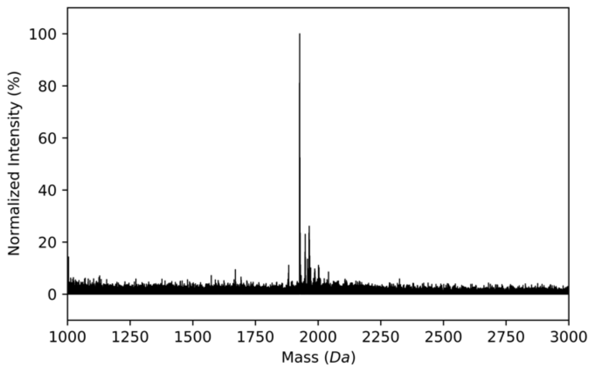


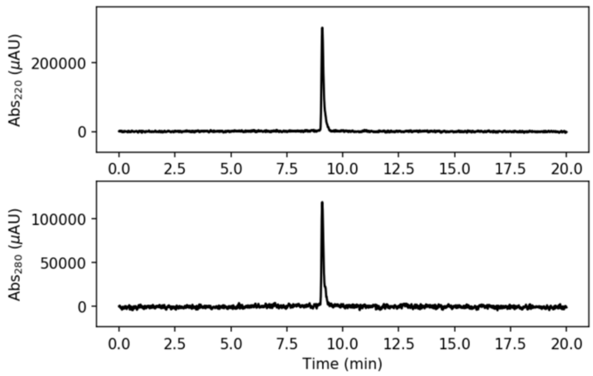


# J-S(L)

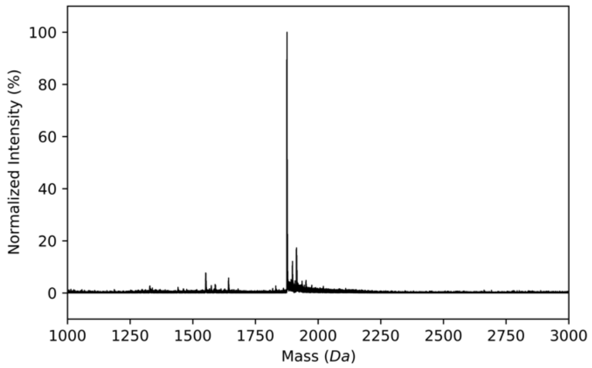


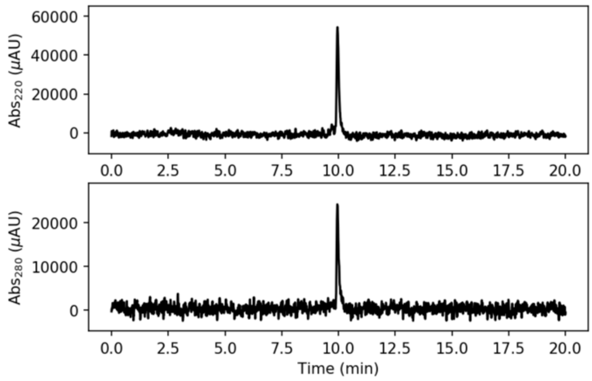

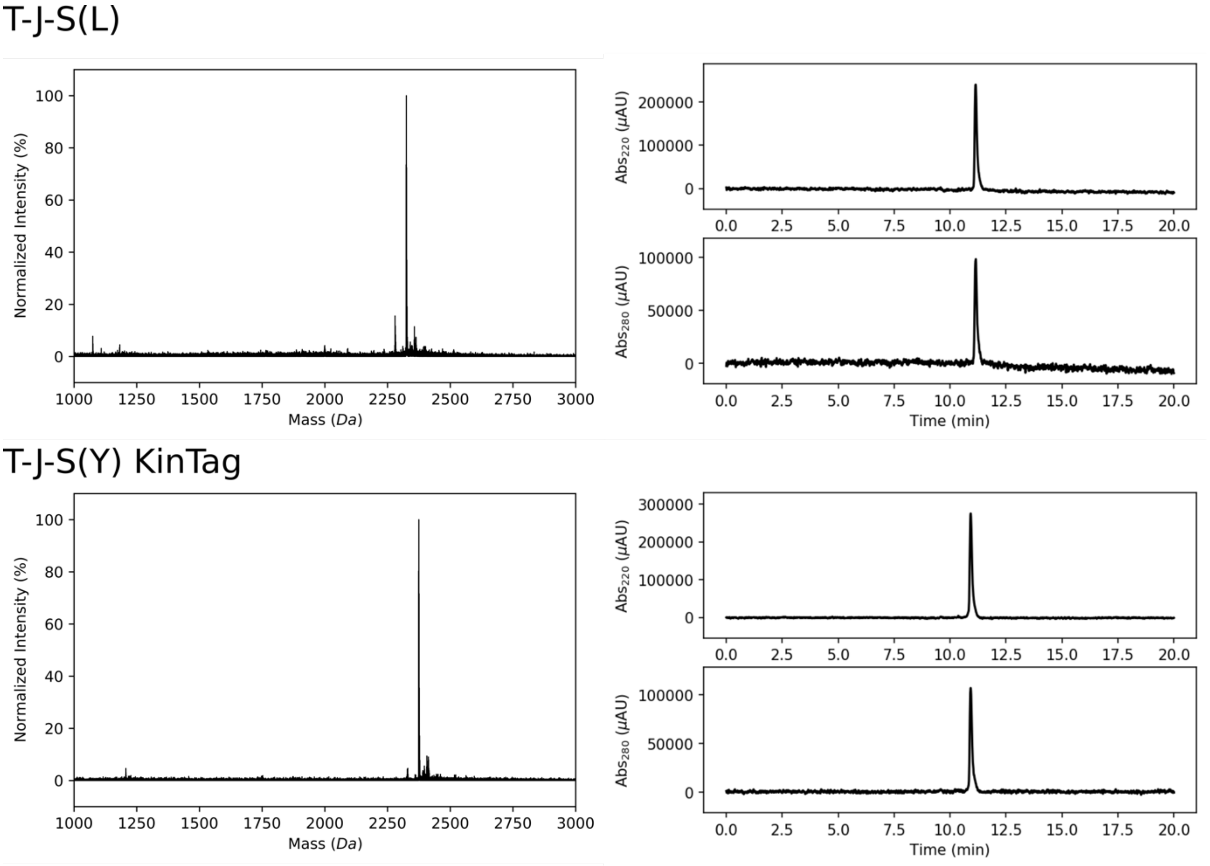


**Supplementary Fig. 1 | Characterisation of synthetic peptides.** Representative MALDI spectra (left) and analytical HPLC traces monitoring absorbance at 220 (top) and 280 nm (bottom) (right) of discussed peptides as shown in table S1.

| **Supplementary Table 2 \| X-ray data collection and refinement** | |
|---|---|
| *Data collection* | |
| Data set | KLC1$^{TPR}$:KinTag |
| Beamline | I03 (DLS) |
| Wavelength (Å) | 0.97625 |
| Resolution range (Å)<br>Highest res. bin (Å) | 111.32-2.85<br>(2.90-2.85) |
| Space group | $P2_12_12_1$ |
| Cell dimensions (Å) | 101.73, 107.53, 222.65 |
| Unique reflections | 57529<br>(2758) |
| Overall redundancy | 4.5<br>(4.4) |
| Completeness, (%) | 99.3<br>(97.1) |
| $R_{merge,}$ (%) | 11.4<br>(154.3) |
| $R_{pim}$ (I), (%) | 6.3<br>(88.9) |
| CC1/2, (%) | 99.4<br>(36.5) |
| ⟨I/σ(I) ⟩ | 8.0<br>(1.2) |
| Wilson *B* factor (Å$^2$) | 46.7 |
| *Refinement* | |
| PDB code | 6SWU |
| $R_{factor}$ (%)/$R_{free}$(%) | 20.1<br>24.3 |
| # non-H atoms | 13791 |
| rmsd bond lengths (Å) | 0.008 |
| rmsd bond angles (°) | 1.421 |

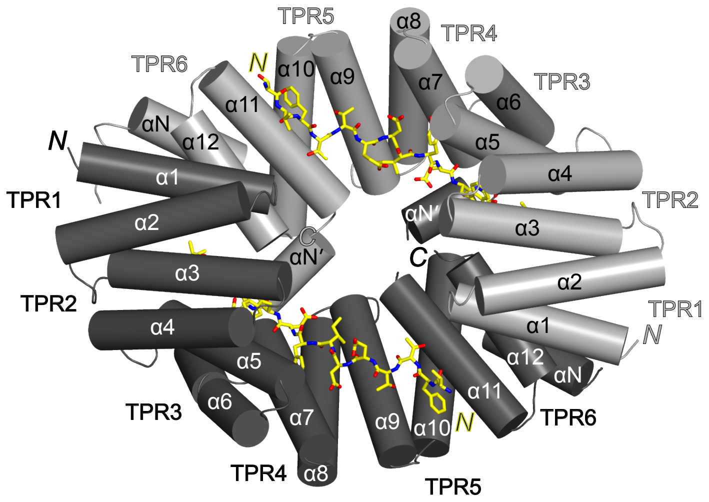


**Supplementary Fig. 2 | KLC1$^{TPR}$:KinTag dimer.** Cartoon representation of the KLC1$^{TPR}$:KinTag head-to-tail dimer viewed approximately down the two-fold axis. KLC1$^{TPR}$ domains are shown in light and dark grey with helices depicted as cylinders. Each KLC$^{TPR}$ domain is formed of six helix-turn-helix TPR repeats (TPR1 α1 - α2, TPR2 α3 – α4, TPR3 α5 – α6, TPR4 α7 -α8, TPR5 α9 – α10, TPR6 α11 – α12) and two additional non-TPR helices (α N and α N') between TPR5 and TPR6. The KinTag peptides are displayed as yellow sticks with nitrogen and oxygen atoms in blue and red, respectively. Colour-coded N and C letters indicate N- and C-termini, respectively. The C-terminus of KinTag peptides is behind α3 helices.

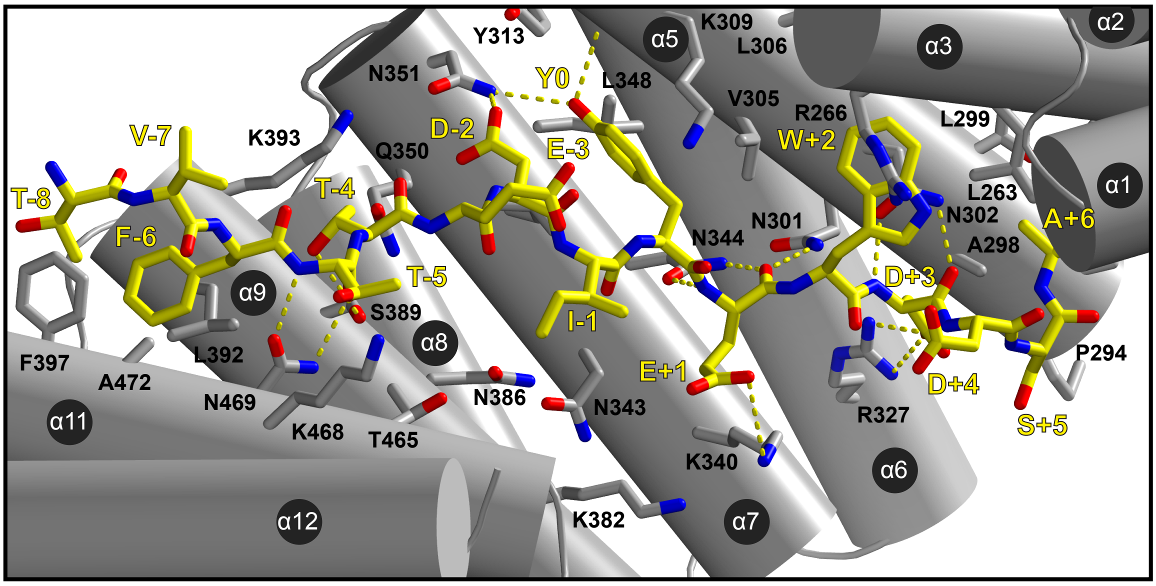


**Supplementary Fig. 3 | KinTag environment and H-bond network.** The KinTag peptide and KLC1$^{TPR}$ residues with which it interacts are shown as sticks in yellow and grey, respectively. Nitrogen and oxygen atoms in blue and red, respectively. Colour-coded labels identify residue numbers. For the KinTag peptide the central tyrosine residue is taken as reference position ($Y^0$). KLC1$^{TPR}$ helices are also labeled. H-bonds and salt bridges are highlighted by broken lines.
